# Mesenchymal-specific Alms1 knockout in mice recapitulates key metabolic features of Alström Syndrome

**DOI:** 10.1101/2023.10.12.562074

**Authors:** Eleanor J. McKay, Ineke Luijten, Xiong Weng, Pablo B. Martinez de Morentin, Elvira De Frutos González, Zhanguo Gao, Mikhail G. Kolonin, Lora K. Heisler, Robert K. Semple

## Abstract

**Background:** Alström Syndrome (AS), a multi-system disease caused by mutations in the *ALMS1* gene, includes obesity with disproportionately severe insulin resistant diabetes, dyslipidemia, and hepatosteatosis. How loss of ALMS1 causes this phenotype is poorly understood, but prior studies have circumstancially implicated impaired adipose tissue expandability. We set out to test this by comparing the metabolic effects of selective *Alms1* knockout in mesenchymal cells including preadipocytes to those of global *Alms1* knockout.

**Methods:** Global *Alms1* knockout (KO) mice were generated by crossing floxed *Alms1* and CAG-Cre mice. A *Pdgfrα*-Cre driver was used to abrogate Alms1 function selectively in mesenchymal stem cells (MSCs) and their descendants, including preadipocytes. We combined metabolic phenotyping of global and *Pdgfrα*+ *Alms1*-KO mice on a 45% fat diet with measurements of body composition and food intake, and histological analysis of metabolic tissues.

**Results:** Global *Alms1* KO caused hyperphagia, obesity, insulin resistance, dyslipidaemia, and fatty liver. *Pdgfrα*-*cre* driven KO of *Alms1* (MSC KO) recapitulated insulin resistance, fatty liver, and dyslipidaemia in both sexes. Other phenotypes were sexually dimorphic: increased fat mass was only present in female *Alms1* MSC KO mice. Hyperphagia was not evident in male *Alms1* MSC KO mice, but was found in MSC KO females, despite no neuronal Pdgfr*α* expression.

**Conclusions:** Mesenchymal deletion of *Alms1* recapitulates the metabolic features of AS, including severe fatty liver. This confirms a key role for *Alms1* in the adipose lineage, where its loss is sufficient to cause systemic metabolic effects and damage to remote organs. AS should be regarded as a *forme fruste* of lipodystrophy. Therapies should prioritise targeting positive energy balance.

## Introduction

Alström syndrome (AS) is a rare recessive disorder caused by biallelic *ALMS1* mutations. Its cardinal features include childhood-onset retinal degeneration, deafness, obesity, diabetes mellitus, and cardiomyopathy ^1^. *ALMS1* encodes a 460kDa protein localised to centrosomes and the basal bodies of primary cilia ^2,3^ but its molecular function remains unknown.

Birthweight in AS is normal, but 98% of affected people develop truncal obesity within one year^4,5^. Insulin resistance (IR) is common, usually severe, and of childhood onset^4,6^. Significantly, IR is disproportionate to adiposity in AS, reportedly five times more severe than in BMI-matched, unaffected controls^7^. It is associated with hypertriglyceridaemia, also often severe, and low HDL cholesterol^8–11^. Severe fatty liver and its sequelae are also common^4–7^. Diabetes, usually of childhood or early adult onset, occurs in 68% of people with AS^4,6,7^. Understanding the pathogenesis of the severe early onset metabolic disease in AS is crucial to improving care for affected people. As the metabolic profile of AS resembles a highly exaggerated form of common obesity-related metabolic syndrome, unpicking the underlying molecular mechanisms may also yield important insights into common disease.

ALMS1 is globally expressed, but the highly lipotoxic subtype of IR seen in AS is reminiscent of the dyslipidaemic severe IR and fatty liver of lipodystrophy. This has led to the hypothesis that the metabolic complications of AS are an exaggerated form of “adipose failure”^6^, albeit without frank anatomical lipodystrophy. Such adipose failure could be attributed to impaired adipocyte differentiation, improper maintenance of the mitotically active adipocyte precursor pool. This is plausible given the role described for primary cilia in early stages of preadipocyte differentiation^12^. A recent murine study, however, has suggested that reactivation of *Alms1* selectively in postmitotic, mature adipocytes substantially mitigates the metabolic consequences of global *Alms1* knockout^13^. While adding to evidence that it is adipose deficiency of *ALMS1* that may explain a raft of crucial metabolic features of AS, this finding was surprising, as it implicated ALMS1 in the function of mature adipocytes, which do not exhibit primary cilia.

We now set out to test whether the metabolic derangement of AS is attributable to loss of *Alms1* in the adipose lineage. To do this we generated both a new global Alms1 knockout (KO) and a new mesenchymal-specific Alms1 KO line by crossing the same floxed Alms1 line with either CAG-cre or Pdgfrα-cre mice^14–16^ and compared the metabolic consequences. While *Pdgfrα-*Cre does target preadipocytes and thus their descendants, endogenous *Pdgfrα* is more accurately viewed as mesenchymal stem cell (MSC) marker^17^. Importantly, however, reporter studies have shown that *Pdgfrα*-Cre driven recombination occurs only at very low levels in liver^16^ and skeletal muscle^15,16^. Thus, the two major peripheral metabolic organs apart from adipose tissue are largely untargeted by the *Pdgfrα* promoter-driven Cre expression. Using this strategy, it is possible to test the role of adipose *Alms1* deficiency in systemic metabolic abnormalities of AS.

## Materials and Methods

### Animal origin and generation

*Alms1*^tm1c(EUCOMM)Hmgu^ (*Alms1*^tm1c^) mice, with loxP sites flanking exon 7 of *Alms1,* were purchased from GenOway. CAG-Cre mice ^18^ were a gift from Dr Matthew Brook and were used to generate global *Alms1* knockout (KO) mice, without potentially confounding Cre expression. *Pdgfrα*-Cre mice ^14^ were purchased from The Jackson Laboratory (Strain #013148) and were used to generate *Alms1* mesenchymal stem cell (MSC) KO mice. Wild type (WT) littermate controls for *Alms1* MSC KO mice expressed *Cre* heterozygously, like their KO littermates. All mice were maintained on a C57/BL6/N background, confirmed by single nucleotide polymorphism profiling (Transnetyx). *Pdgfrα*-Cre x mTom/mGFP mice have been previously described^19^.

*Alms1* expression is normal in *Alms1*^tm1c^ mice, but *Cre* expression excises exon 7 and introduces a nonsense mutation in exon 8 (**Figure S1A**). In the absence of a reliable antibody against murine Alms1, KO was validated by real-time PCR of genomic DNA (Transnetyx) and of cDNA from metabolic tissues studied. PCR amplification of cDNA across exon 7 indicated complete loss of the short exon 7 in *Alms1* global KO mice, and partial loss in MSC KO mice (**Figure S1B**). Quantitative real time PCR (Supplementary Methods) of cDNA targeting an amplicon including the *Alms1* exon 6-7 junction in inguinal white adipose tissue (iWAT) and liver corroborated total loss of *Alms1* in global KO mice and partial loss of *Alms1* in MSC-specific KO adipose tissue (**Figure S1C,D**). Differences in liver in MSC-KO mice were not significant, although a trend to a mild reduction was seen in females (**Figure S1E,F**).

### Animal husbandry

*Alms1* Global- and MSC- KO mice were group-housed in individually ventilated cages (IVCs) at the Biological Research Facility at the University of Edinburgh where a 12-hour light/dark cycle (lights on 0700; off 1900) and controlled temperature/humidity (19-21°C/50%) were maintained. Prior to 6 weeks old, mice had *ad libitum* access to standard chow (CRM, Special Diet Service). All experimental protocols were approved by the University of Edinburgh Biological Science Services in compliance with the UK Home Office Scientific Procedure (Animals) Act 1983.

### In vivo metabolic phenotyping

At 6 weeks of age, mice were transferred from standard chow to a 45% fat diet (D12451, Research Diets; caloric value 4.73kcal/g), with water and diet available *ad libitum*. Body mass was measured weekly from 6 to 24 weeks of age. Body composition was measured by time-domain nuclear magnetic resonance (tdNMR) using the Bruker Minispec Live Mice Analyzer LF50 ^20^. Areas Under the Curve (AUC) for body mass and composition were calculated for each animal. Body length was measured using a digital calliper from nose-anus under anaesthesia at 24 weeks prior to cardiac puncture. Mice were single housed from 14 to 18 weeks of age, with food mass measured at the start and end of the 4-week period, with the difference taken as a measure for food intake. For linear regression between food intake and lean mass, the lean mass at 14 weeks of age was used. Metabolic efficiency was calculated by dividing the energy associated with change in body mass by food intake for the same period, expressed as a percentage. Change in body mass was defined as change in lean mass (kJ) plus change in fat mass (kJ), with fat and lean mass assigned energy content of 39kJ/g and 5kJ/g respectively^21^.

Unfasted blood was collected from the tail vein at 10 and 19 weeks of age at 1100 for glucose and insulin assay. Glucose concentration was determined from the tail vein using the AccuChek Performa Nano [Roche, Switzerland]. Insulin was assayed by electrochemiluminescence immunoassay at the Medical Research Council Metabolic Diseases Unit (MRC-MDU) Mouse Biochemical Assay Laboratory at the University of Cambridge (Supplementary Methods). At 20 and 22 weeks of age, an Insulin Tolerance Test (ITT) and oral Glucose Tolerance Test (oGTT) were performed, with lean mass measured by tdNMR one day beforehand. Mice were fasted from 0700 until testing at 1100. Glucose was measured in tail vein blood using the AccuChek Performa Nano. For ITT, insulin was delivered by intra-peritoneal injection, with 1.75 mIU Humulin S/g lean mass for females, and 2.5 mIU/g lean mass for males used. For oGTT, 4 mg glucose/g lean mass was delivered by oral gavage as 25% D-glucose in saline. Blood samples were collected in EDTA-coated capillary tubes and placed on ice before spinning at 4°C for 10 mins at 2000g and separating plasma. Plasma was stored at -80°C and thawed for biochemical analysis. Area of glycaemia curve (AOC) for oGTT and ITT were calculated for each individual animal, with t0 glycaemia as baseline^22^. For oGTT, the area above t0 glycaemia, and for ITT, the area below t0 glycaemia were quantified.

At 24 weeks, a terminal bleed was undertaken by cardiac puncture under general anaesthesia and tissues were harvested immediately post mortem, weighed, and sectioned before fixing of tissues in 4% paraformaldehyde or snap freezing in liquid nitrogen. Plasma biochemical profiling including cytokine assays were performed in the MRC-MDU (Supplementary Methods).

### Histological studies

Paraformaldehyde-fixed tissues were paraffin-embedded and 5μm sections cut before hematoxylin and eosin (H&E) and picrosirius red (PSR) staining using standard procedures^23,24^. Sudan Black B (SBB) staining was performed as described previously^25^, but due to the fragility of adipose sections, wiping to remove excess stain was avoided. Immunohistochemical (IHC) staining for p21 (ab188224, 1:500, Abcam, UK), p16 (AHP1488, 1:500, BioRad, USA) and Lamin-B1 (ab16048, 1:100, Abcam, UK) was undertaken using a Leica BOND III automated immunostainer robot at room temperature together with staining with haematoxylin and 3,3′-Diaminobenzidine (H-DAB) using the Bond Polymer Refine Detection Kit. Liver and adipose histological slides were imaged using a Ziess Axioscan.Z1 with Zen2.6 software. Images at 20x magnification were analysed in QuPath v0.3.1^26^. SBB, and DAB staining in adipose and liver sections was quantified using an automated self-defined pixel classifier. PSR-staining was quantified as detailed in Supplementary Methods. Detection of brain mTomato (anti-RFP, rabbit, 1:1000, 600-401-379, Rockland Immunochemicals, USA) and mGFP (anti-GFP, chicken, 1:1000, ab13970, Abcam, UK) in Pdgfrα:Cre x mTom/mGFP mice by dual immunohistochemistry used methods adapted from those previously described^27^, and is detailed further in Supplementary Methods.

### Blinding and Statistical analysis

Experimenter and analyser were blinded to genotype where possible, including for *in vivo* animal work, dissections, RNA extraction and histological processing and analysis. cDNA synthesis and qPCR were unblinded to enable generation of pooled control reverse transcriptase negative reactions for different experimental groups. Unblinding of data was automated in Microsoft Excel template to prevent accidental memorisation of genotypes. Statistical analysis was performed in GraphPad Prism 9.2.0. A normal distribution was assumed for all data, as the small numbers required by best practice in animal research preclude reliable testing for normality. Student’s t-test was used to compare data sets pairwise, and ANOVA for comparison of more than 2 groups. Linear regression was used to compare differences between two experimental groups in their relationship between two dependent variables (e.g. food intake and lean mass). Bonferroni correction was applied when multiple t-tests were performed on the same graph, as when plotting global and MSC-specific KO data. Šídák’s multiple comparisons test was applied following ANOVA when the same animals were studied at more than one time point. Tukey’s multiple comparison test was applied following ANOVA to compare values among multiple groups. Statistical analysis was performed on raw data except for insulin values which were log transformed first.

## Results

### Generation of global and mesenchymal stem cell-specific Alms1 knockout mice

To test the hypothesis that all or most of the lipotoxic IR in AS is explained by Alms1 deficiency in the MSC/preadipocyte/adipocyte lineage, novel mouse lines with either global or mesenchymal stem cell (MSC-) specific knockout (KO) of *Alms1* were generated. This was achieved by crossing mice with floxed *Alms1* exon 7 with others expressing *Cre* either constitutively (CAG-Cre) for global KO, or in *Pdgfrα*-expressing cells only^14–16^ for MSC-specific KO.

### MSC-specific Alms1 deficiency has sexually dimorphic effects on appetite

To ensure vigorous recruitment of adipose tissue, mice were fed a 45% HFD *ad libitum* from 6 weeks of age. In agreement with previously published models, both male and female global *Alms1* KO mice had higher body mass than wild-type (WT) controls (**Figure 1A-D**). Female *Alms1* MSC KO mice recapitulated this increased body mass (**Figure 1A,B**), but male *Alms1* MSC KO mice showed only a similar trend (**Figure 1C,D**). The increased body mass of female global- and MSC- KO mice and male global KO mice was accounted for by increased fat mass (**Figure 1E-H**), with lean mass unchanged (**Figure S2A-D**).

**Figure 1.**
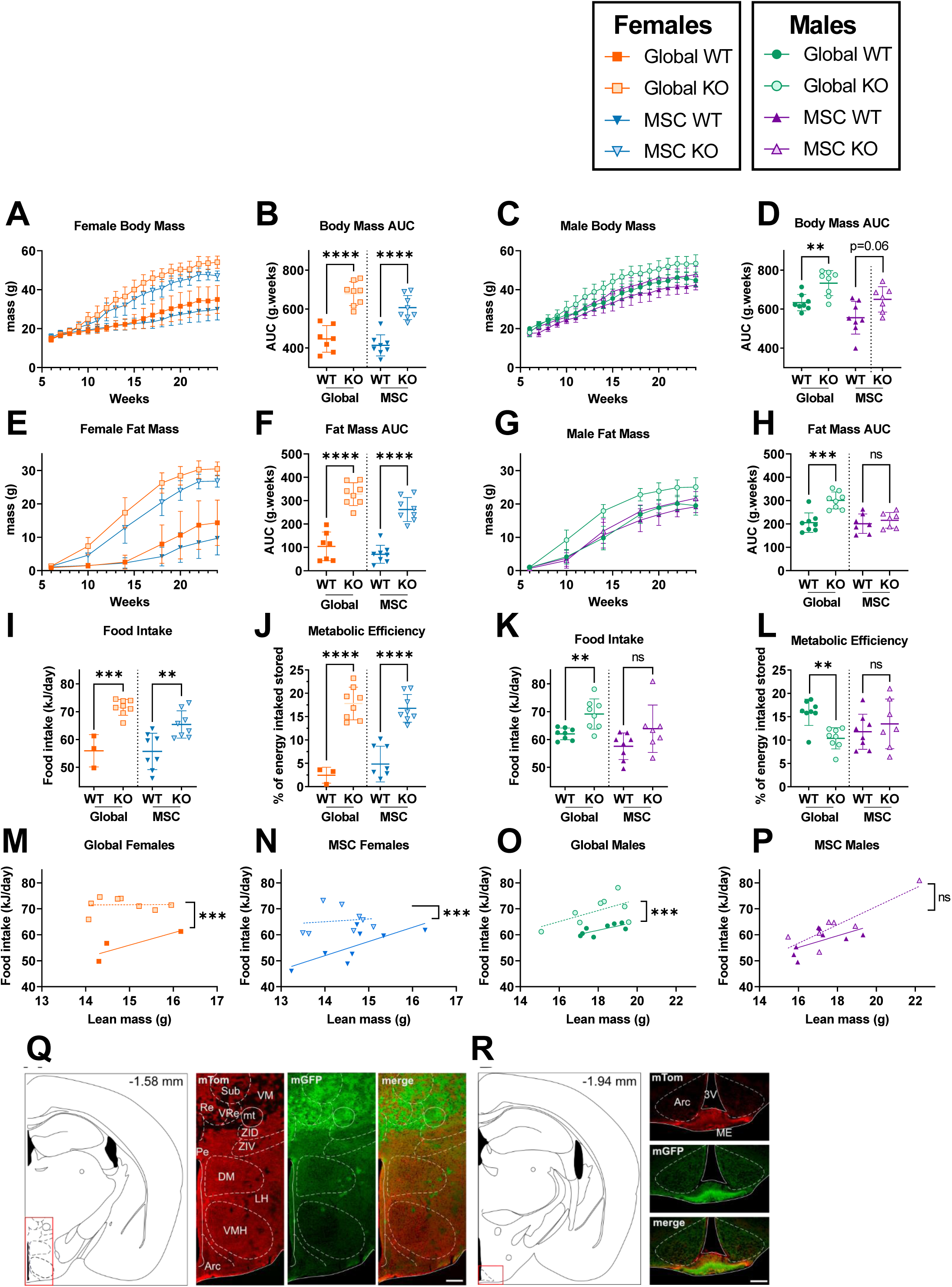
Female, but not male, MSC-specific *Alms1* knockout mice recapitulate the obesity and hyperphagia of global *Alms1* knockout. Longitudinal analysis of body mass (A,C) and fat mass (E,G) assessed by td-NMR for female (A,E) and male (C,G) mice on high fat diet. (I-P) Food intake and metabolic efficiency of animals measured from 14-18 weeks of age. (M-P) Food intake plotted against lean mass at 14 weeks of age. Comparisons of global WT and KO and MSC WT and KO were performed with identical design at different times, reflected in the dotted line separating comparisons. Longitudinal series (A,C,E,G) plot mean ± sd for each time point. Data points in (B,D,F,H-P) represent individual animals with bars in (B,D,F,H-L) representing mean ± sd, and lines in linear regression graphs (M-P) representing lines of best fit. Comparison between WT and KO in (A-L) was performed using an unpaired two-tailed Student’s t-test with Bonferroni correction. Comparison between lines of best fit was performed by simple linear regression, with square brackets showing comparison of y intercepts. No significant change was seen between gradients. ** denotes p<0.01, *** denotes p<0.001 and **** denotes p<0.0001. For females N = 7, 8, 8 and 8 for global WT, global KO, MSC WT and MSC KO respectively, except in food intake studies when many global WT females shred the diet, resulting in N=3. For males N = 8, 8, 7 and 7 for global WT, global KO, MSC WT and MSC KO respectively. AUC = area under curve. (Q,R) Illustration of coronal plane and representative microphotographs of brain sections from female Pdgfrα-Cre x mTom/mGFP mice showing expression of mTom, mGFP and merged images at bregma levels (Q) -1.58; (R) -1.94. Arc: arcuate hypothalamic nucleus; DM: dorsomedial hypothalamic nucleus; LH: Lateral hypothalamic area; mt: mammillothalamic tract; Pe: periventricular hypothalamic nucleus; Sub: supratrigeminal nucleus; VM: ventromedial thalamic nucleus; VRe: ventral reuniens thalamic nucleus; ZID: zona incerta, dorsal part; ZIV: zona incerta, ventral part; 3V: Third ventricle. Scale bar 200 µm for Q and 100 µm for R.

Food intake was measured from 14-18 weeks of age while the mice were single-caged. As expected, both female and male global *Alms1* KO mice were hyperphagic (**Figure 1I,K**), and increased food intake remained significant when lean body mass was taken into account (**Figure 1M,O**). Unexpectedly, given neuronal sparing of *Pdgfrα*-Cre, female *Alms1* MSC KO mice, but not males, recapitulated the hyperphagia of global *Alms1* loss (**Figure 1N,P**). Increased metabolic efficiency, defined as the percentage of energy intake that is stored as lean or fat mass, was also seen in female *Alms1* global- and MSC- KO mice (**Figure 1J,L**). In contrast, male global *Alms1* KO mice showed decreased metabolic efficiency, with no difference between male MSC KO and controls (**Figure 1J,L**).

In view of the surprisingly hyperphagia of female mice with *Alms1* KO limited to *Pdgfrα*-expressing lineages, we next examined brains from *Alms1* WT, *Pdgfrα*-Cre mice harbouring an mGFP/mTomato reporter cassette. This expresses mTomato by default, switching to mGFP on Cre exposure. Evidence of recombination and thus *Pdgfrα-*Cre expression at some point in development was detected in several brain regions. No obvious sexual dimorphism was noted. Extremely low levels of recombination were seen in regions classically associated with feeding behaviour such as the dorsomedial, ventromedial, lateral or arcuate nucleus of the hypothalamus (**Figure 1Q,R**). Similar to a previous report^28^, a population of *Pdgfrα*-Cre recombined cells was visualised in the midline of the median eminence (**Figure 1R**). Robust recombination was also seen in the reuniens thalamic nucleus (**Figure 1Q**), the rostral periolivary region and intermediate nucleus of the lateral lemincus (**Figure S1E**), anterodorsal thalamic nucleus (**Figure S1F**), superior cerebellar peduncle (**Figure S1G**) and area postrema (**Figure S1H**). Low level co-expression of mTomato and mGFP in some regions could be explained by transient developmental expression of *Pdgfrα*, while scattered cells lacking both mTomato and mGFP are consistent with observations made during generation of the mTom/mGFP reporter model^29^. These findings are in keeping with the expected expression of *Pdgfrα* in oligodendrocyte precursor cells (OPCs) but not neurones.

### MSC-specific Alms1 deficiency is sufficient to induce dyslipidaemic insulin resistance

To assess the presence of the systemic metabolic complications of AS, we started by determining non fasting blood glucose and insulin concentrations in mice at 10 and 19 weeks of age. Female global *Alms1* KO mice were hyperglycaemic at both time points, and MSC KO mice only at 19 weeks (**Figure 2A**), with concomitant plasma insulin concentrations markedly increased at all times where hyperglycaemia was observed (**Figure 2B**). Male mice showed no hyperglycaemia at 10 or 19 weeks (**Figure 2C**), but at both time-points, global *Alms1* KO mice were severely hyperinsulinaemic, with insulin concentrations around 10-fold higher than controls. Normoglycaemic hyperinsulinaemia was recapitulated in male *Alms1* MSC KO mice at 19 weeks of age (**Figure 2D**).

**Figure 2.**
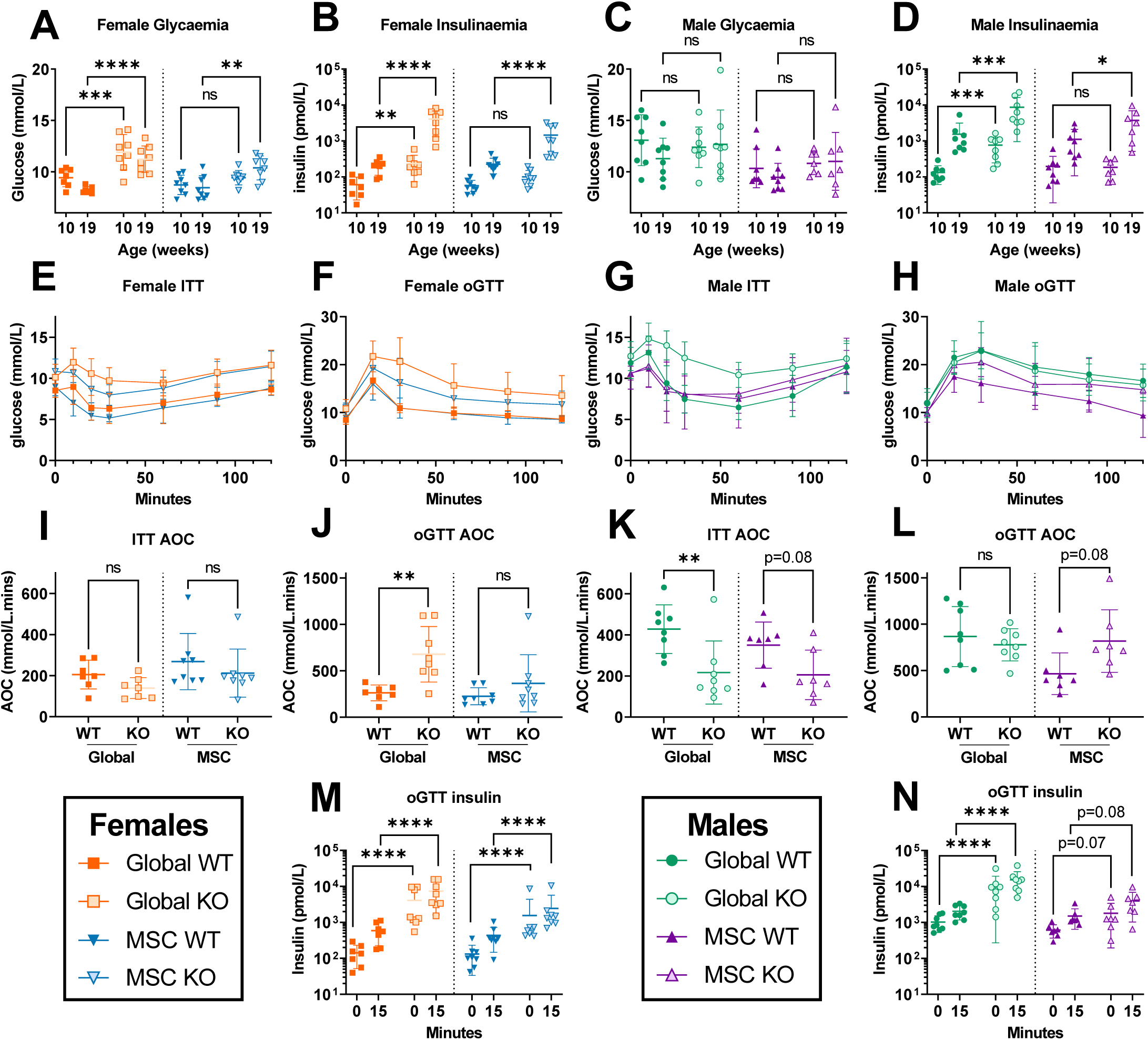
Mesenchymal stem cell-specific *Alms1* knockout recapitulate the insulin resistance of global *Alms1* loss. (A-D) Daytime non-fasted blood glucose and insulin concentrations at 10 and 19 weeks of age. (E,G) insulin tolerance testing (ITT) and (F,H) oral glucose tolerance testing (oGTT). Global WT/KO and MSC WT/KO experiments were performed with identical design at different times, reflected in the dotted line separating comparisons in (A-D, I-N). oGTT and ITT graphs (E-H) show mean ± sd for each time point. All other graphs (A-D, I-N) plot data points representing individual animals, with bars representing mean ± sd. Comparisons of log10 oGTT insulin values (B,D,M,N) and glucose concentration (A,C) were performed by two-way ANOVA with Šídák’s multiple comparisons test. Comparison of all other data (I-L) used an unpaired two-tailed Student’s t-test with Bonferroni correction. * denotes p<0.05, ** denotes p<0.01, *** denotes p<0.001 and **** denotes p<0.0001. For females N = 7, 8, 8 and 8 for global WT, global KO, MSC WT and MSC KO respectively. For males N = 8, 8, 7 and 7 for global WT, global KO, MSC WT and MSC KO respectively. AOC = area of the curve; ns = not significant.

Insulin and oral glucose tolerance tests (ITT and oGTT) were then undertaken at 20 and 22 weeks of age respectively to gain a more dynamic assessment of glucose homeostasis. These showed no significant reduction in hypoglycaemic response to insulin in female *Alms1* global- or MSC- KO mice (**Figure 2E,I**) but increased glycaemic excursion after oral glucose in global KO mice, with a trend only in MSC KO animals (**Figure 2F,J**). Both global- and MSC- KO female mice showed severe hyperinsulinaemia (**Figure 1M**). Male global *Alms1* KO mice showed a reduced hypoglycaemic response to insulin (**Figure 2G,K**), while MSC KO in males produced a trend towards reduction (**Figure 2K**). Global *Alms1* KO males showed unaltered glycaemic excursion after oral glucose (**Figure 2H,L**), but severe hyperinsulinaemia (**Figure 2N**), and MSC KO mice showed a trend towards increased glucose excursion and insulinaemia on oGTT (**Figure 2L,N**). Collectively these findings demonstrate severe IR in global KO mice of both sexes, which is largely recapitulated in MSC KO mice. Hyperglycaemia, however, was only prominent in female global KO mice, being only weakly echoed in female MSC KO mice.

The IR in AS is reported to be of a strikingly dyslipidaemic subtype^7^. To confirm this, we also determined lipid profiles in the KO models. At 24 weeks, global *Alms1* KO mice were indeed strikingly hyperlipidaemic (Table 1), albeit with a different pattern to human metabolic dyslipidaemia, which characteristically features elevated serum triglyceride and suppressed HDL cholesterol. The global *Alms1* KO mice mice had higher serum total, HDL and LDL cholesterol in both sexes, but higher triglyceride in males only (**Table 1**). Male MSC KO mice showed no changes in lipid profile, while female MSC KO mice showed hypercholesterolaemia only. Circulating free fatty acid (FFA) concentrations showed no genotype-related differences.

**Table 1:**
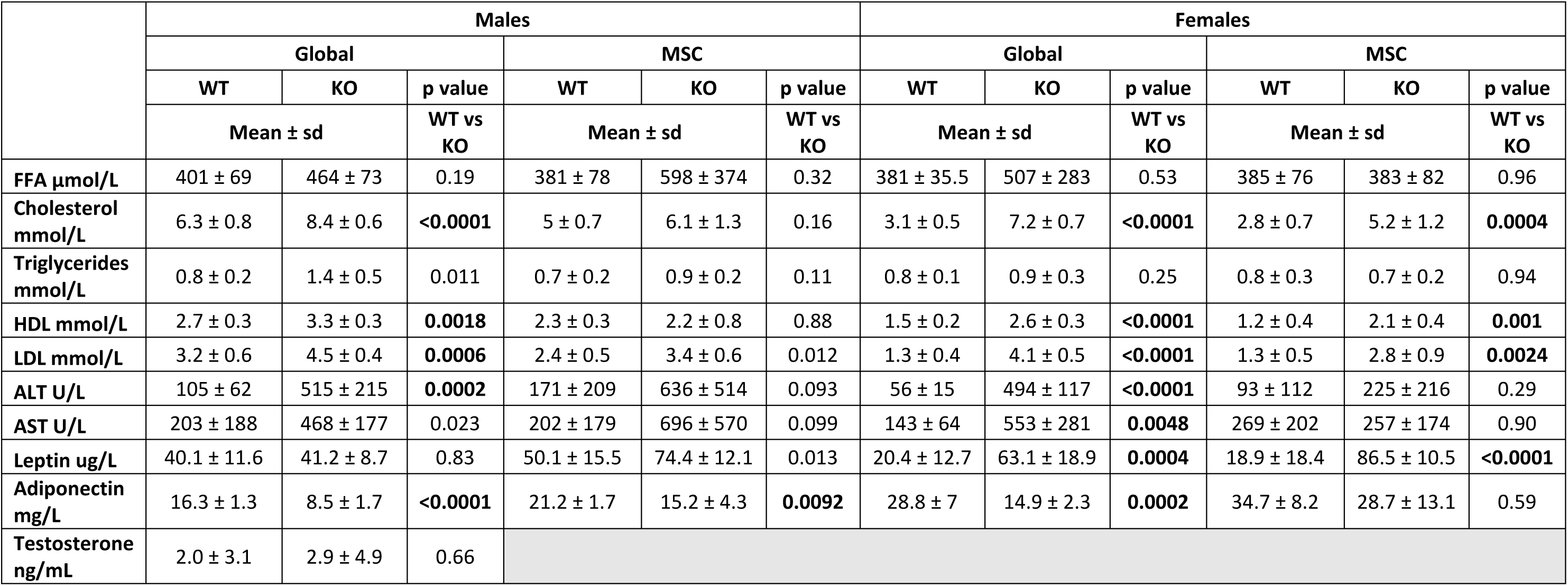
Serum biochemical profiles of global and MSC-specific *Alms1* KO mice. Data are shown for high fat fed mice at 24 weeks of age. Global WT/KO and MSC WT/KO experiments were performed with identical design at different times. WT/KO comparison used an unpaired two-tailed Student’s t-test with Bonferroni correction. For females N = 7, 8, 8 and 8 for global WT, global KO, MSC WT and MSC KO respectively. For males N = 8, 8, 7 and 7 for global WT, global KO, MSC WT and MSC KO respectively. P values below 0.005 (given 10 comparisons) are shown in bold.

### Effects of MSC-specific Alms1 deficiency on adipose cellularity are also sexually dimorphic

Consistent with the increased adiposity of female *Alms1* global- and MSC- KO mice, serum leptin concentration was increased in both. Male global *Alms1* KO mice appeared to show no difference in serum leptin concentration despite differences in fat mass, but this is likely artefactual, as leptin concentrations reached the upper limit of detection in the assay used. In keeping with this, a trend in increased serum leptin concentration was seen in male *Alms1* MSC KO mice, for which sufficient plasma was obtained for repeat assay of diluted samples. Serum adiponectin concentrations were reduced in male and female global *Alms1* KO mice, and in male but not female MSC KO mice. As adiponectin is suppressed by testosterone, and given common male hypogonadism in AS, serum testosterone was assayed in male global *Alms1* KO mice, but was unaltered compared to controls.

Having confirmed hyperlipidaemic IR in global *Alms1* KO, as in human AS, and to a significant extent in MSC KO mice, we then assessed mass and cellularity of adipose depots. At 24 weeks old, masses of inguinal and gonadal white adipose tissue depots (iWAT and gWAT respectively) and liver were greater in female global and MSC KO mice than controls (**Figure 3A-D**). Interscapular brown adipose tissue (iBAT) mass showed a trend towards increase in global KO females, and a significant increase in MSC KO female mice. WAT depot masses showed a divergent pattern in males. iWAT mass was increased, while gWAT mass was decreased in global KOs, suggesting WAT redistribution. Neither change was recapitulated in MSC KO mice (**Figure 3E,F**). No differences in iBAT mass were seen in males. When serum leptin concentrations were plotted against fat mass, no genotype-related difference was seen for either sex (**Figure 3I-L**).

**Figure 3.**
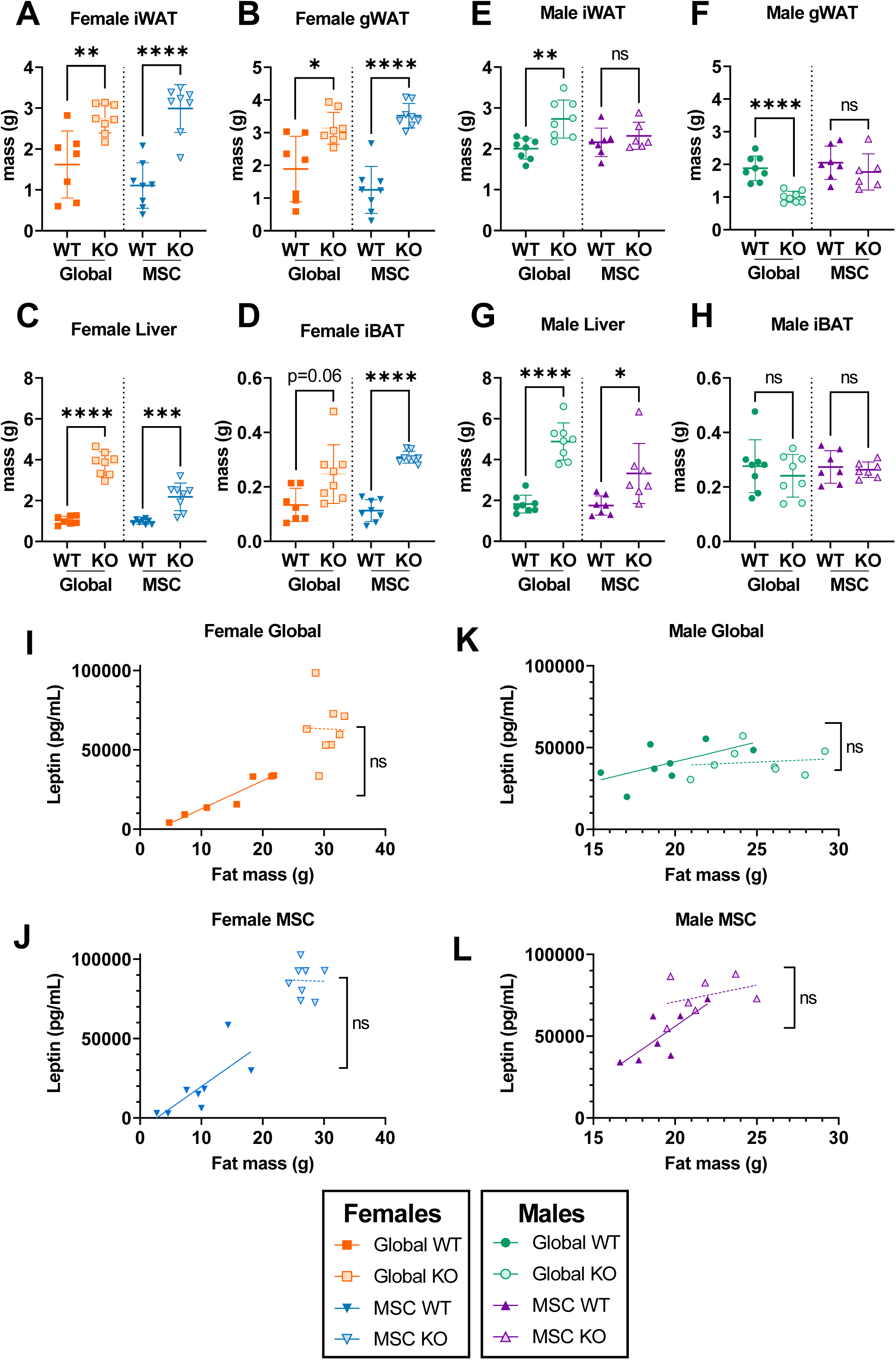
Mesenchymal stem cell-specific *Alms1* knockout mice show hepatomegaly despite the absence of Cre-driven loss of Alms1 in hepatocytes, while fat pad mass is sexually dimorphic. (A-H) Mass of inguinal, and gonadal white adipose tissue (iWAT and gWAT respectively), liver and interscapular brown adipose tissue (iBAT) of male and female mice at 24 weeks of age. (I-L) Linear regression of serum leptin concentration and lean mass of male and female mice at 24 weeks of age. Each data point represents an individual animal, with bars in (A-H) representing mean ± sd lines and lines in (I-L) representing lines of best fit. Comparison between WT and KO groups in (A-H) used an unpaired two-tailed Student’s t-test with Bonferroni correction. Comparison between lines of best fit in (I-L) was performed by simple linear regression, with square brackets showing comparison of y intercepts. No significant change was seen between gradients. * denotes p<0.05, ** denotes p<0.01, *** denotes p<0.001 and **** denotes p<0.0001. For females N = 7, 8, 8 and 8 for global WT, global KO, MSC WT and MSC KO respectively. For males N = 8, 8, 7 and 7 for global WT, global KO, MSC WT and MSC KO respectively. ns = not significant.

Cross sectional area of adipocytes was next measured for iWAT and gWAT (**Figure 4C,D & S3A,B**). Female iWAT and gWAT showed increased adipocyte size for both *Alms1* global- and MSC- KO animals (**Figure 4I,J & S3C,D**) while iWAT of male global *Alms1* KO mice showed increased adipocyte size that was not recapitulated by MSC KO (**Figure 4L,M**). Contrary to all other WAT depots investigated, male gWAT showed reduced adipocyte size in global *Alms1* KO (**Figure S3F**), but this was not seen in MSC KO mice (**Figure S3G**). No change in calculated adipocyte numbers was seen for any depot except iWAT of female MSC KO mice (**Figure 4K,N & S3E,H**).

**Figure 4.**
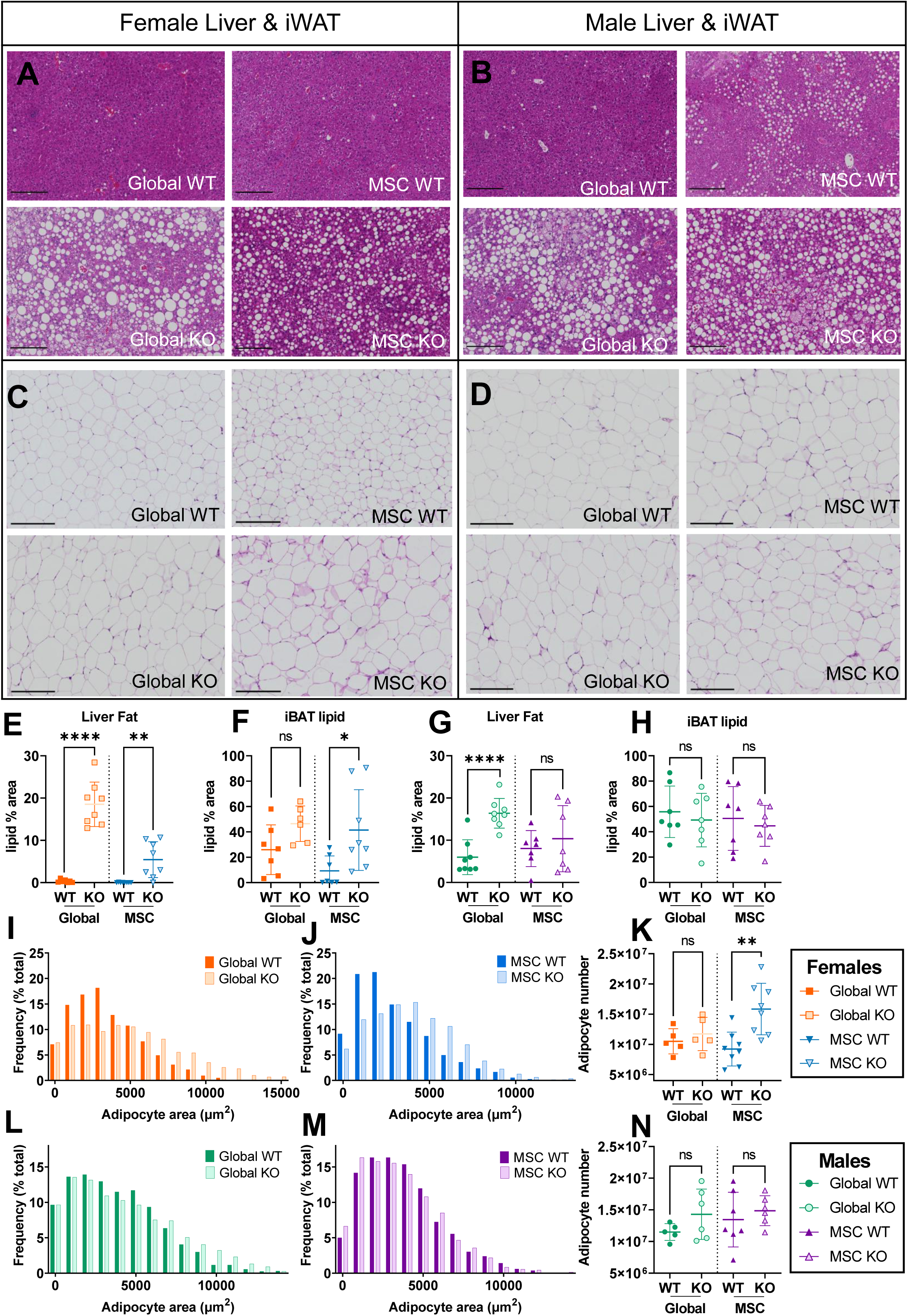
Adipocyte hypertrophy and increased liver fat in both global and mesenchymal stem cell-specific *Alms1* knockout mice at 24 weeks of age. Representative images of haematoxylin and eosin (H&E) stained liver and inguinal white adipose tissue (iWAT) sections from (A,C) female and (B,D) male mice. Scale bars 200μm. Quantification of lipid content of liver (E,G) and interscapular brown adipose tissue (iBAT) (F,H) sections. (I,J,L,M) Size distribution of cross sectional area of adipocytes in iWAT, represented in bins of 1000μm^2^ from comparisons of WT and global or conditional KO mice in separate experiments. (K,N) Total number of adipocytes in iWAT of each animal. Each data point in (E-H,K,N) represents an individual animal, with bars representing mean ± sd. Comparison between WT and KO in (E-H,K,N) used an unpaired two-tailed Student’s t-test with Bonferroni correction. * denotes p<0.05, ** denotes p<0.01 and **** denotes p<0.0001. (E-H) For females N = 7, 8, 8 and 8 for global WT, global KO, MSC WT and MSC KO respectively. For males N = 8, 8, 7 and 7 for global WT, global KO, MSC WT and MSC KO respectively. (I-N) For females N = 5, 5, 8 and 8 for global WT, global KO, MSC WT and MSC KO respectively. For males N = 5, 6, 7 and 7 for global WT, global KO, MSC WT and MSC KO respectively. ns = not significant.

### MSC-specific Alms1 deficiency has lipotoxic effects on remote tissues with intact Alms1 expression

Adipocyte hypertrophy and dyslipidaemic IR are consistent with failure of adipose tissue to launch an adequate hyperplastic response to positive energy balance. Such “adipose failure” characteristically leads to ectopic accumulation of lipids in remote organs, including liver and thermogenic brown adipose tissue. Liver mass was markedly increased in both global- and MSC- KO mice (**Figure 3G**), with pale macroscopic appearances of the KO livers (**Figure S4A,B**) and histological evidence of macrovesicular steatosis (**Figure 4A,B,E,G**), although in male MSC KO mice the difference in histological lipid quantification did not reach statistical significance. Serum alanine transaminase and aspartate transaminase concentrations, indices of hepatocellular damage, were increased in both sexes of *Alms1* global-, but not MSC- KO mice (**Table 1**).

Lipid deposition in iBAT, assessed by quantification of the proportion of the cross-sectional area accounted for by lipid droplets, tended to be increased in female global KO and was significantly increased in MSC KO (**Figure 4F & S4C**) but unchanged in male global and MSC KO mice (**Figure 4H & S4D**). Collectively, these findings support the presence of ectopic lipid accumulation or “lipotoxicity” in global and MSC-specific KO animals, but with some sexual dimorphism, as for other traits assessed.

### Alms1 knockout induces systematic meta-inflammation without adipose fibrosis or senescence

Widespread fibrosis of many organs, including liver and WAT, is reported in human AS, and is the target in an ongoing clinical trial^30^, but it is unclear whether this mediates metabolic derangement and organ dysfunction, or is simply an epiphenomenon. We undertook picrosirius red (PSR) staining of histological sections of liver, iWAT, and gWAT of global and MSC-specific *Alms1* KO mice to assess for fibrosis, but detected no significant changes in the extent of staining (**Figure S5A-F**), although a trend towards increase was seen in gWAT of female global KO mice.

Adipocellular senescence, both cell autonomous and paracrine, has also been invoked as a significant contributor of obesity complications with age, and senolytic therapies have emerged as encouraging candidates to mitigate metabolic disease progression^31^. Given evidence of adipocyte hypertrophy, and expression of ALMS1 in the centrosome and mitotic spindle, we thus also undertook histological analysis of cellular senescence markers in iWAT and gWAT of global *Alms1* KO mice. However no difference in staining with Sudan Black B (SBB), which binds lipofuscin^32^ (**Figure S5G,H**), nor immunohistochemical staining p16, p21 and Lamin B1 (**Figure S5I-N**) was seen. We also determined serum concentrations of a panel of cytokines including some constituents of the senescence-associated secretory phenotype (SASP) (**Table 2**). Female global KO mice showed marked elevation of several cytokines, including core SASP components IL-6, and CXCL-1, and variable SASP components IL-10 and TNFα, however IL-1β, one of the most reliable SASP markers, was unchanged. Male global KO mice showed a less striking profile of cytokines, although IL-6, CXCL1 and IL-10 were elevated. MSC KO mice broadly failed to reproduce the pattern of increases in cytokines seen in global KO animals, though female MSC KO mice did show increases in TNFα and IL-6 of borderline significance. These findings support the presence of marked meta-inflammation in global Alms1 KO animals, and suggest that Alms1 expression in non *Pdgfrα* lineages such as immune cells may be involved in this. However, the question of whether there is a significant burden of senescent cells in AS remains to be resolved definitively.

**Table 2:**
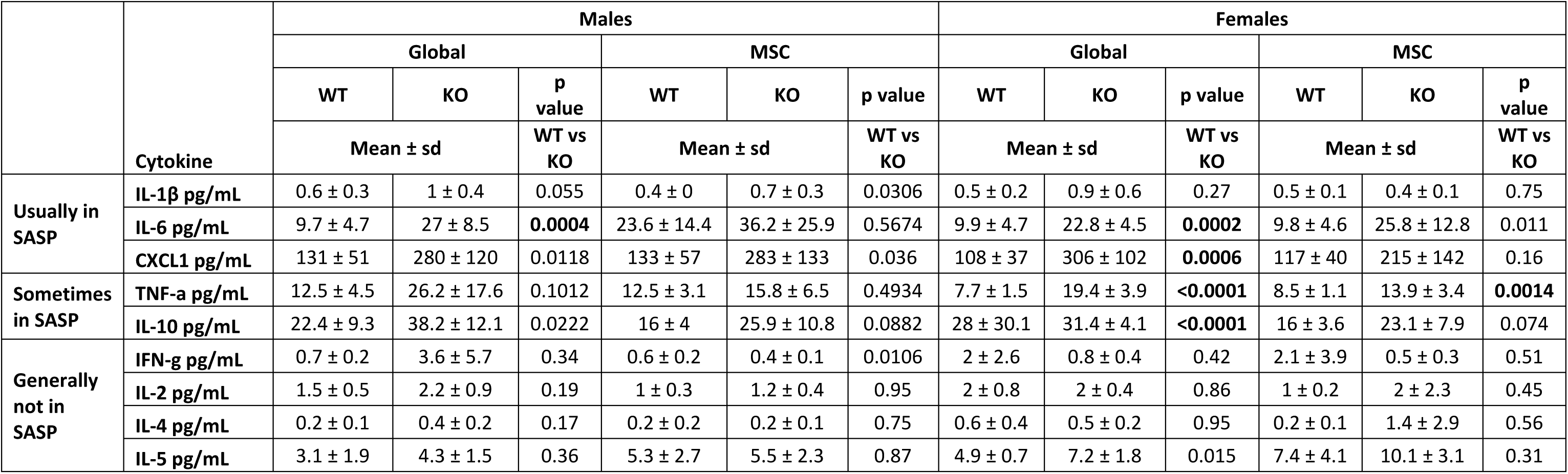
Serum cytokine analysis in global and MSC-specific *Alms1* KO mice. Data are shown for high fat fed mice at 24 weeks of age. Global WT/KO and MSC WT/KO experiments were performed with identical design at different times. WT/KO comparison used an unpaired two-tailed Student’s t-test with Bonferroni correction. For females N = 7, 8, 8 and 8 for global WT, global KO, MSC WT and MSC KO respectively. For males N = 8, 8, 7 and 7 for global WT, global KO, MSC WT and MSC KO respectively. P values below 0.006 (given 9 comparisons) are shown in bold.

## Discussion

The adipose expandability hypothesis holds that when the capacity of adipose tissue to expand by hyperplasia in response to positive energy balance is exceeded, adipocyte hypertrophy and necrosis ensue, leading to dyslipidaemic IR^33,34^. Adipocyte hypertrophy has been shown in adipose tissue of patients with AS ^13^, and the severely dyslipidaemic IR and fatty liver of AS strongly resemble the metabolic profile of lipodystrophies, extreme examples of “adipose failure”. We have thus suggested that loss of *Alms1* in the adipocyte lineage may be sufficient to explain many or all of the metabolic complications of AS. We predicted that *Pdgfra-*Cre driven *Alms1* knockout would induce not only severe IR, but also severe fatty liver disease, despite intact hepatocyte *Alms1* expression. We also sought to determine whether *Pdgfra*-driven *Alms1* knockout would modify the inflammation and fibrosis characteristic of the liver and adipose pathology of human AS^6^.

The metabolic phenotype of the global *Alms1* KO mice we report accords with other reported global KO models^35–37^. In line with best practice, we studied mice of both sexes, and this revealed some sexual dimorphism, however. The greater metabolic derangement and obesity induced by *Alms1* KO in female compared to male mice, although not previously discussed, is consistent with the greater body weight increase reported in females than males in three prior *Alms1* KO mouse models^35–38^. Some of this sexual dimorphism may relate to differences in susceptibility of male and female WT C57BL/6 mice to diet-induced-obesity. Females are usually resistant to this more than males, potentially making pathologically increased susceptibility easier to discern^39,40^. No sexual dimorphism of metabolic traits has been reported in human AS, however the rarity of AS and lack of dedicated studies of sex differences to date means that some sexual dimorphism of metabolic complications remains possible.

In keeping with our hypothesis that *Alms1* deficiency in the MSC/preadipocyte/adipocyte lineage accounts for the severe IR of AS, female *Alms1* MSC KO mice recapitulate the metabolic phenotype of global KO almost completely. Male *Alms1* MSC KO mice, although not exhibiting the obesity of *Alms1* global KO mice, also showed IR. The hepatic steatosis in female *Alms1* MSC KO is notable given intact hepatocyte *Alms1* expression, confirming that fatty liver in AS does not require liver autonomous Alms1 deficiency. On the other hand, the degree of fatty liver is smaller in MSC KO mice than global KO mice, and indeed in males there is no significant change. Moreover the increased serum ALT and AST of global KO mice is not recapitulated by *Alms1* MSC KO mice. This may implicate non *Pdgfrα*-Cre lineages in the hepatocellular damage that complicates fatty liver, but caution is warranted as global and MSC KO animals were not studied in parallel. The greater fatty liver in female than male MSC KO mice may also reflect the hyperphagia of female but not male mice, which would increase the caloric load on adipose tissue.

Increased mass of iWAT and gWAT depots in female global and MSC KO mice, without an increase in adipocyte numbers per depot, is consistent with constrained recruitment of new preadipocytes. Increased iBAT lipid content, like fatty liver, is evidence of iBAT “whitening”, most likely due to lipid overspill from WAT. Neither adipose fibrosis nor overt histological evidence of adipocyte senescence were seen, though raised circulating cytokines indicate that metinflammation, and possibly senescence, may be at play systemically. This evidence of systemic inflammation was much less pronounced in MSC KO mice, suggesting, as in liver, that Alms1 loss in non *Pdgfrα* lineages may play a role in some of the sequalae of adipose failure. Decoupling of fibrosis from adipose failure and dyslipidaemic IR provides evidence that fibrosis itself is not a prerequisite for these metabolic complications, and argues against anti-fibrotic strategies to treat metabolic complications of AS.

Observations of male KO adipose tissue were more nuanced. Although a trend towards hypertrophy was seen in iWAT, global *Alms1* KO males had decreased gWAT mass without hypertrophy, in agreement with findings in the *Alms1^foz/foz^*model^41^. iWAT showed no such reduction in our study or others ^41^. Neither iWAT nor gWAT mass were changed in male *Alms1* MSC KO mice, suggesting that the adipose redistribution observed in males is not accounted for by MSC-derived cells. Lineage tracing studies have shown that iWAT shows high levels of recombination with a *Pdgfrα*-Cre driver, while some non-recombined adipocytes persist in gWAT^19^. It is not clear which sex these experiments were performed in, but this raises the possibility of a methodological basis for our findings.

Generating *Adipoq*-Cre driven adipocyte-specific *Alms1* KO will be an important future experiment to establish whether *Alms1* loss is causing adipose tissue failure due to effects on preadipocytes or mature adipocytes. This is a significant question given surprising prior data suggesting that restoration of *Alms1* function only in mature adipocytes of global *Alms1* KO mice ameliorates the adverse metabolic phenotype^13^.

An unexpected finding in this study was that the hyperphagia of global *Alms1* KO was also seen in female - but not male – MSC KO mice. Hyperphagia in *Alms1^foz/foz^* mice precedes increased fat mass^36,42^, while a pair-feeding study showed that hyperphagia was a significant contributor to obesity and IR^42^. Typically, such appetitive phenotypes are neuronally mediated, hypothalamic neuronal cilia have been implicated in appetite control^43^, and *Alms1^foz/foz^* mice have fewer ciliated cells in the hypothalamus ^44^. We did not observe evidence for *Pdgfrα*-driven recombination in hypothalamic appetitive nuclei, however. Recombination was seen instead in the median eminence and elsewhere in a pattern consistent with oligodendrocyte precursor cells (OPCs), as expected given widespread use of *Pdgfrα*-Cre^ER^ to manipulate OPC gene expression. OPCs are thus strong candidate mediators of hyperphagia in response to *Alms1* KO. There is in fact growing evidence for a role for OPCs and their descendants in appetite control^28,45,46^. Mechanisms invoked include secondary loss of leptin receptors in the arcuate nucleus^45^, and remodelling of perineuronal nets in response to nutritional fluxes in the median eminence^28^. Impaired differentiation of OPCs into oligodendrocytes due to impaired ciliary transmission of nutrition-related signals, leading to lower perineuronal net formation and less neuronal satiety signalling is a hypothesis worthy of examination in Alms1 KO mice. OPCs are highly sensitive to sex hormones, and sexual dimorphism in OPC number, proliferation and migration is well established^47^. We speculate that this may explain the sexually dimorphic hyperphagia of MSC KO mice.

In conclusion, our findings are consistent with a key role for Alms1 in adipocyte hyperplasia and thus buffering of chronic positive energy balance. Loss of Alms1 limited to MSC-derived cells including preadipocytes is sufficient to cause relative adipose failure on a high fat diet, with secondary fatty liver and dyslipidaemia despite intact hepatocyte *Alms1* expression. Surprisingly, our findings also raise the possibility that the hyperphagia of AS may be attributable to loss of *Alms1* in oligodendrocyte precursor cells as well as, or instead of, neurones. Importantly, we do not see evidence of fibrosis nor overt cellular senescence in either adipose tissue or liver, despite severe systemic metabolic derangement. This argues against targeting these as therapeutic strategies in AS. Efforts to reverse positive energy balance are likely to be a higher yield approach to offloading failing adipose tissue and improving health.

## Author Contributions

**EJM:** conceptualisation, methodology, formal analysis, investigation, writing - original draft, writing - review & editing, visualization. **IL**: supervision, writing - review & editing. **XW**: investigation. **PMM:** methodology, investigation, visualisation. **EDFG:** investigation. **ZG** & **MGK**: resources. **LKH:** resources, supervision, writing - review & editing. **RKS:** conceptualisation, methodology, resources, writing - original draft, writing - review & editing, supervision, funding acquisition.

