## Supplementary Information for "Mesenchymal-specific Alms1 knockout in mice recapitulates key metabolic features of Alström Syndrome"

### Supplementary Table

| Metabolite | Reagent manufacturer | Product code |
|---|---|---|
| Triglycerides | Siemens Healthcare | DF69A |
| AST | Siemens Healthcare | DF41A |
| ALT | Siemens Healthcare | DF143 |
| Cholesterol | Siemens Healthcare | DF27 |
| HDL | Randox | CH2849 |
| FFA | Roche | Sigma 11383175001 |
| Testosterone | Demeditec | DEV9911 |
| Insulin & Leptin | MesoScale Discovery | K15124C-3 |
| Adiponectin | MesoScale Discovery | K152BYC-2 |
| Inflammatory cytokines | MesoScale Discovery | K15048D-2 |

**Supplementary Table 1. Reagents used for serum biochemical asssays.**

### Supplementary Figures

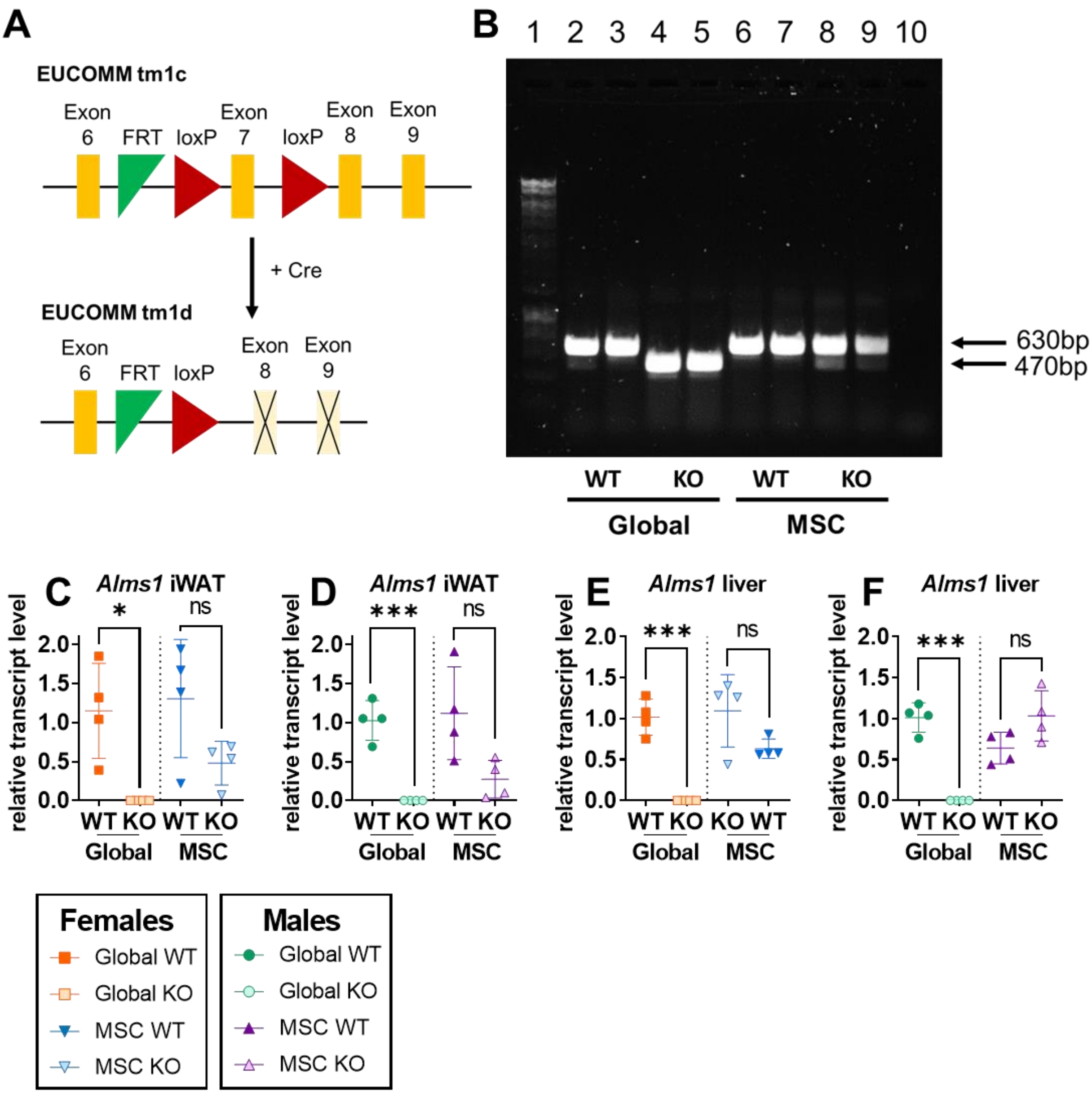


**Figure S1. Validation of global and MSC-specific *Alms1* KO mice.** (**A**) Schematic of *Alms1*$^{tm1c}$ and *Alms1*$^{tm1d}$ alleles, both generated from the EUCOMM *Alms1*$^{tm1c}$ allele. (**B**) Liver cDNA gel electrophoresis following PCR amplification across area including exon 7 which is 160bp long. Lane 1: DNA ladder; lanes 2&3: *Alms1* global WT cDNA amplicon; lanes 4&5: *Alms1* global KO cDNA amplicon; lanes 6&7: *Alms1* MSC WT cDNA amplicon; lanes 8&9: *Alms1* MSC KO cDNA amplicon; lane 10, non-template control. (**C-F**) qPCR validation of *Alms1* loss in global and MSC-specific Alms1 KO mice using a Taqman probe spanning the exon 6-7 junction of *Alms1*, normalised to *Gapdh*. Global WT/KO and MSC WT/KO experiments were performed with identical design at different times, reflected in the dotted line separating the two cohorts. N = 4/group. Data represent individual animals with bars representing mean ± sd. Comparison between WT and KO in (**C-F**) used an unpaired two-tailed Student's t-test with Bonferroni correction. iWAT = inguinal white adipose tissue

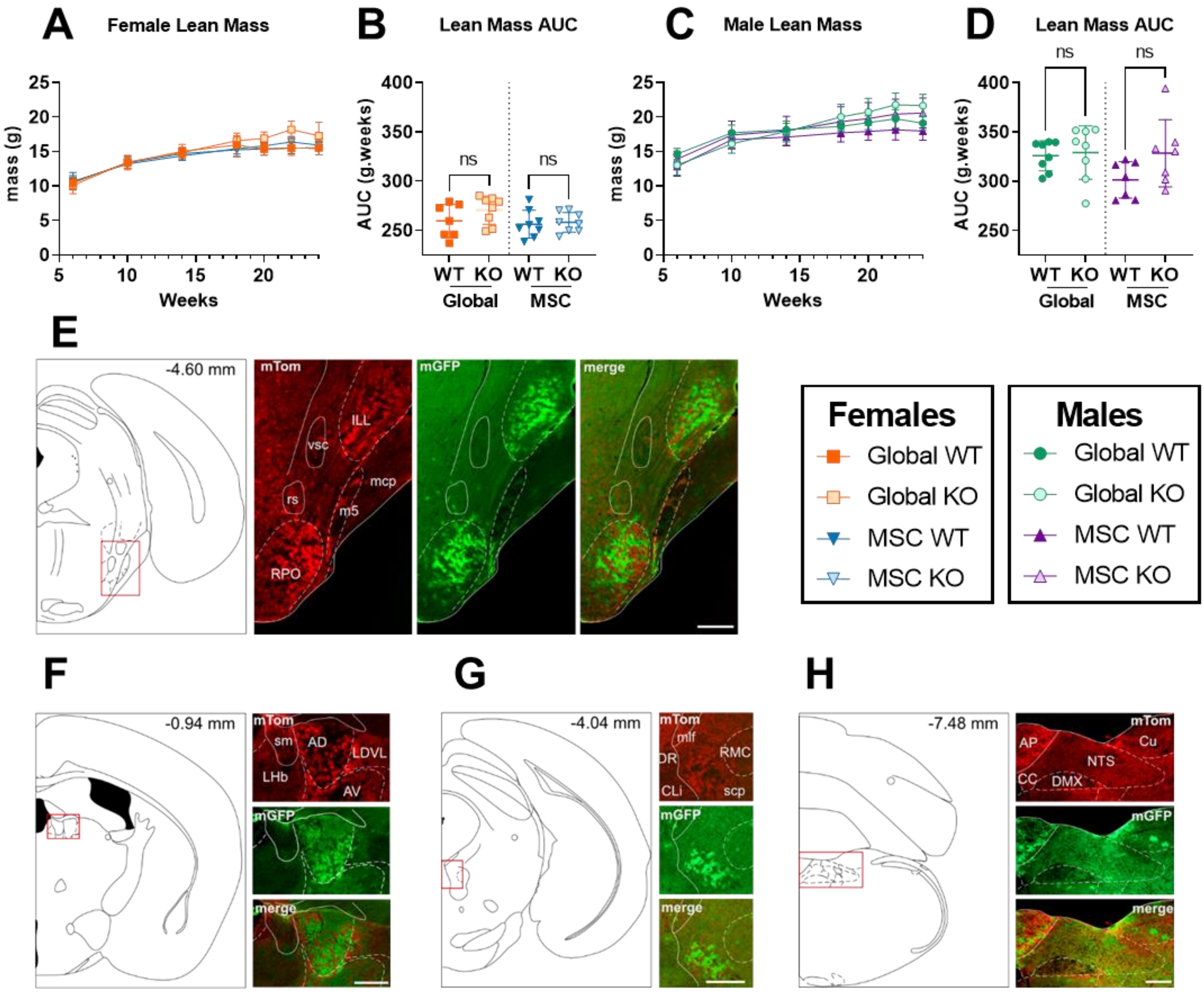


**Figure S2. Female, but not male, MSC-specific *Alms1* knockout mice recapitulate the obesity and hyperphagia of global *Alms1* knockout.** (**A-D**) Longitudinal analysis of lean mass assessed by td-NMR for female (**A**) and male (**C**) mice on high fat diet. For females N = 7, 8, 8 and 8 for global WT, global KO, MSC WT and MSC KO respectively. For males N = 8, 8, 7 and 7 for global WT, global KO, MSC WT and MSC KO respectively. AUC = area under curve WT/Global KO and WT/MSC KO experiments were performed with identical design at different times, reflected in the dotted line separating the comparisons. Longitudinal series plot mean ± sd for each time point. Data points in (**B,D**) represent individual animals with bars representing mean ± sd. Comparison between WT and KO in (**B,D**) was performed using an unpaired two-tailed Student's t-test with Bonferroni correction. **(E-H)** Illustration of coronal plane and representative microphotographs of brain sections from female Pdgfrα-Cre x mTom/mGFP mice showing expression of mTom, mGFP and merged images at bregma levels **(E)** -4.6; (**F**) -0.94; (**G**) -4.04; and (**H**) -7.48. AD: anterodorsal thalamic nucleus; sm: stria medullaris of the thalamus; AP: area postrema; AV: anteroventral thalamic nucleus; CC: central canal; CLi: caudal linear nucleus of the raphe; Cu: cuneate nucleus; DMX: dorsal nucleus of the vagus nerve; DR: dorsal raphe; LDVL: laterodorsal thalamic nucleus, ventrolateral part; LHb: lateral habenular nucleus; mlf: medial longitudinal fasciculus; NTS: nucleus of the solitary tract; RMC: red nucleus, magnocellular part; scp: superior cerebellar peduncle. Scale bar 200 µm.

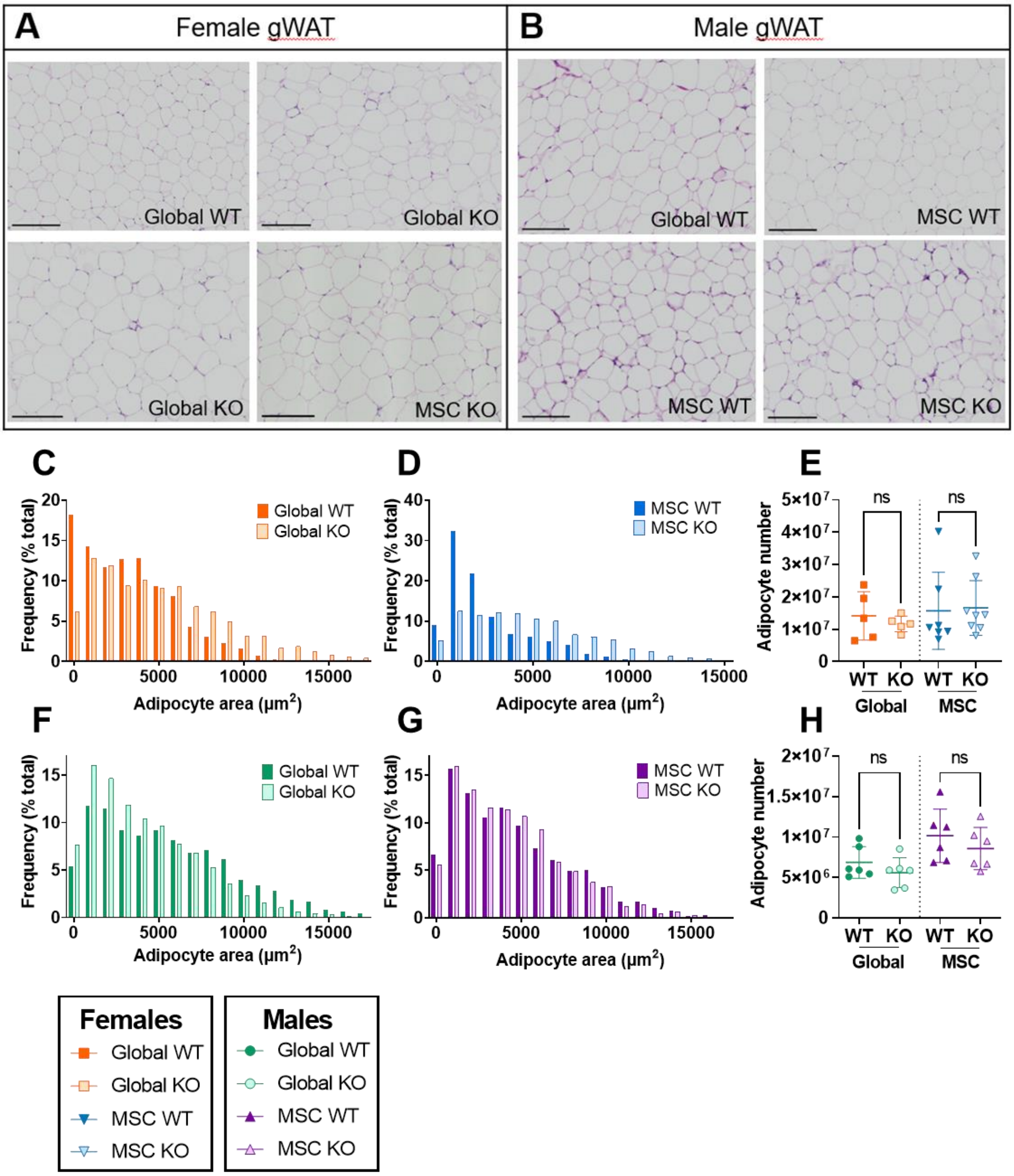


**Figure S3. Histological evaluation of gonadal white adipose tissue of global and mesenchymal stem cell-specific *Alms1* knockout mice.** (**A,B**) Representative microscopic images of haematoxylin and eosin-stained gonadal white adipose tissue (gWAT) sections from female and male WT, global or mesenchymal stem cell (MSC) knockout (KO) mice at 24 weeks of age on high fat diet. Scale bars 200μm. (**C,D,F,G**) Size distribution of cross sectional area of adipocytes in gWAT, represented in bins of 1000μm$^2$. (**E,H**) Calculation of the total number of adipocytes in the gWAT depot of each animal. Each data point represents one animal, with bars representing mean ± sd. Comparison between WT and KO groups in used an unpaired two-tailed Student's t-test with Bonferroni correction for multiple testing. (**C-H**) For females N = 5, 5, 8 and 8 for global WT, global KO, MSC WT and MSC KO respectively. For males N = 6, 6, 7 and 7 for global WT, global KO, MSC WT and MSC KO respectively.

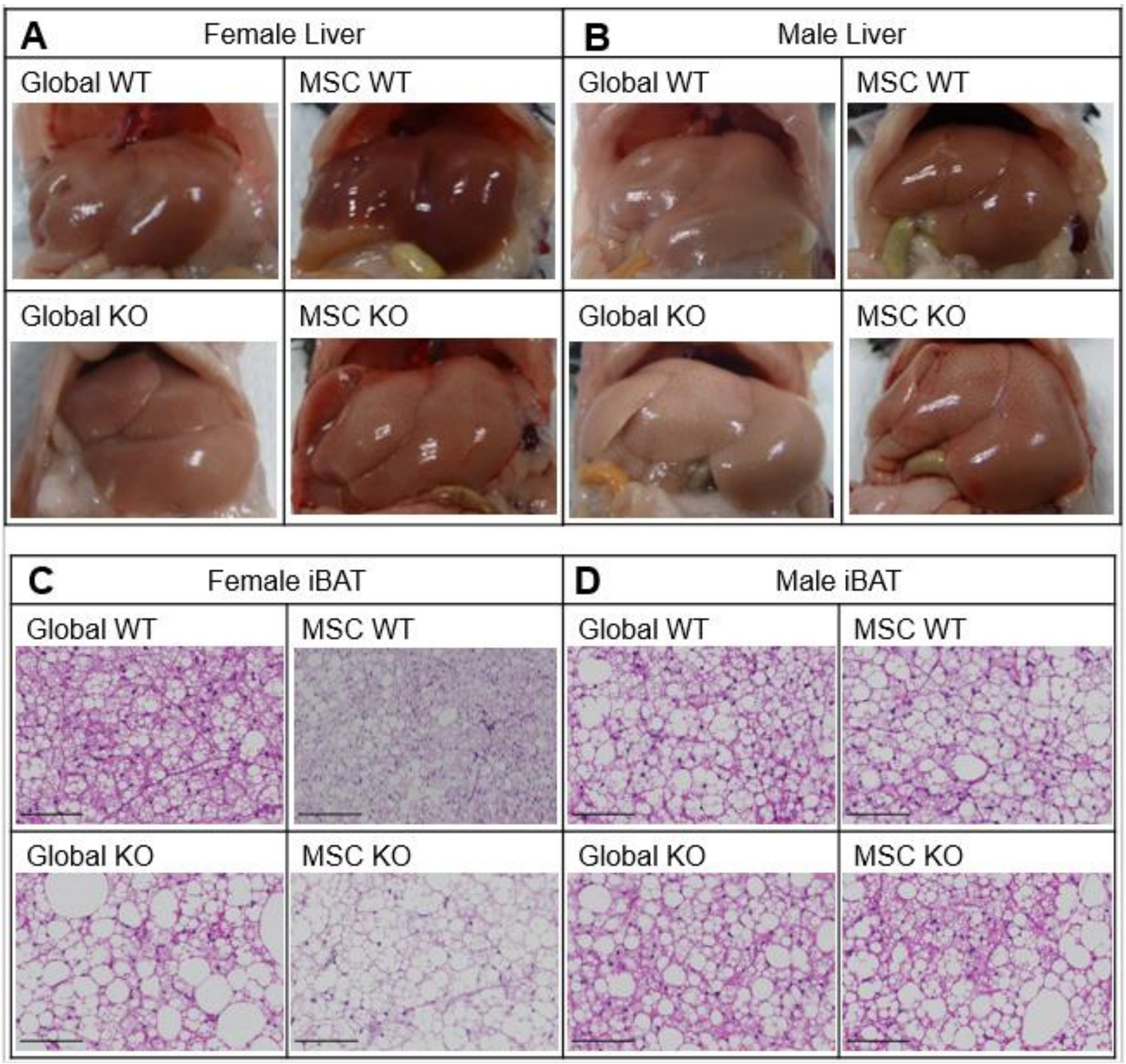


**Figure S4. Histological evaluation of liver and interscapular brown adipose tissue of global and MSC-specific *Alms1* knockout mice.** Representative macroscopic images of liver (**A,B**) and microscopic images of haematoxylin and eosin stained interscapular brown adipose tissue (iBAT) (**C,D**) from WT, global and mesenchymal stem cell (MSC) *Alms1* knockout (KO) mice at 24 weeks of age. Scale bars are 200µm.

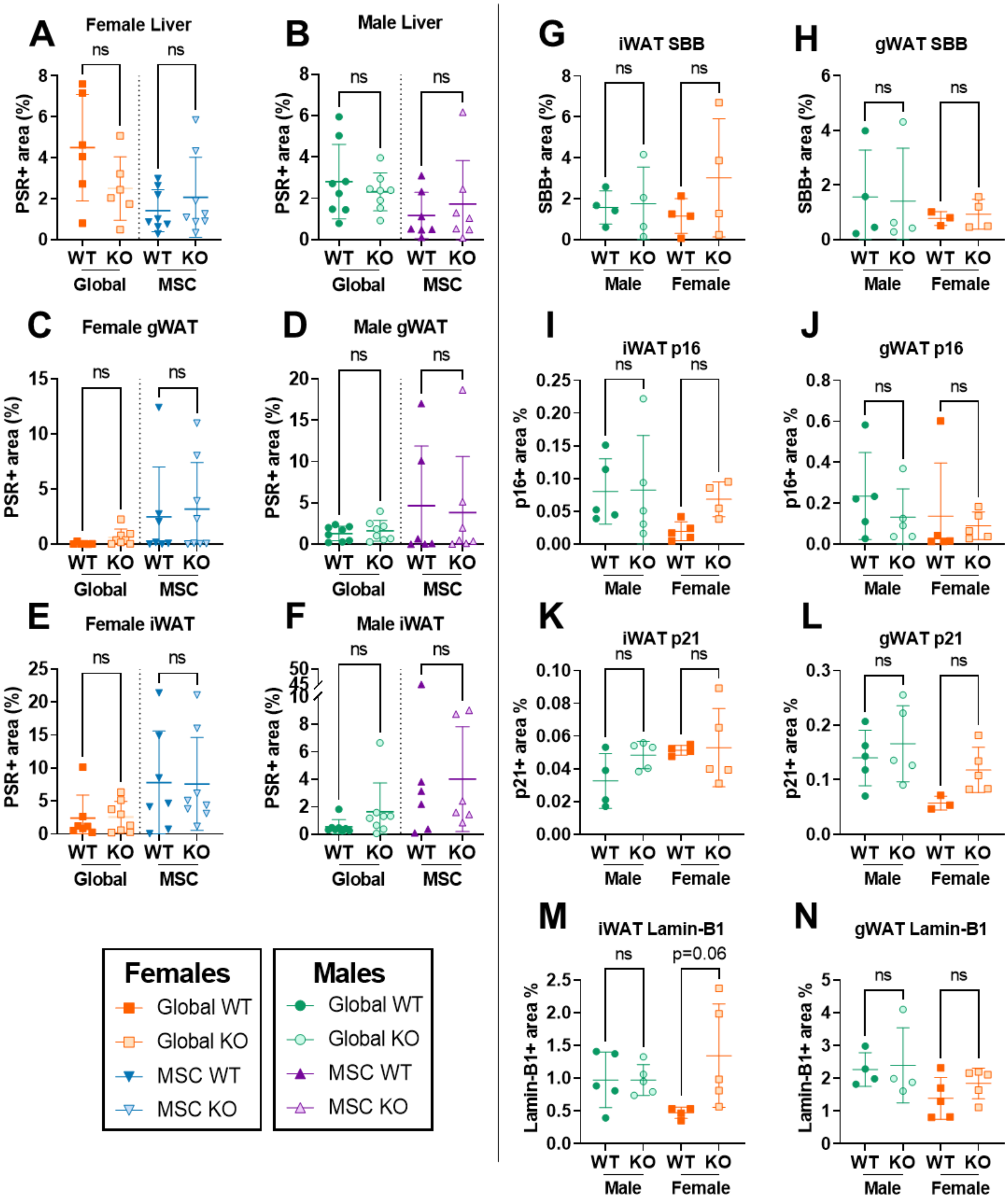


**Figure S5. Histological evaluation of metabolic tissues from global and mesenchymal stem cell-specific *Alms1* knockout mice.** Quantification of picrosirius red (PSR) fibrosis staining in liver (**A,B**), gonadal white adipose tissue (gWAT) (**C,D**) and inguinal WAT (iWAT) (**E,F**) in 24 week old male and female global and mesenchymal stem cell (MSC) knockout (KO) mice. (**G-N**) Quantification of senescence markers in gWAT and iWAT: (**G,H**) Sudan Black B staining for lipofuscin. (**I-N**) Immunohistochemical staining for (**I,J**) p16 (**K,L**) p21 and (**M,N**) LaminB1. Each data point represents one animal, with bars representing mean ± sd.

Comparison between groups in (**A-F**) used an unpaired two-tailed Student's t-test with Bonferroni correction. Comparisons in (**G-N**) used two way ANOVA with Tukey's multiple comparison test. * denotes p<0.05, ** denotes p<0.01 and **** denotes p<0.0001. (**A-F**) For females N = 7, 8, 8 and 8 for global WT, global KO, MSC WT and MSC KO respectively. For males N = 8, 8, 7 and 7 for global WT, global KO, MSC WT and MSC KO respectively. (**G,H**) N = 4/group. (**I-N**) N = 5/group.

### Supplementary Methods

*Histology in Pdgfrα:Cre x mTom/mGFP brain*

For the detection of mTom and mGFP brains were dissected, post-fixed in formalin and cryoprotected with 30% sucrose at 4°C before coronal sectioning in 5 series at 25 µm using a freezing microtome (8000, Bright Instruments, UK). Sections were stored in anti-freeze solution at 4°C until processing. Briefly, sections were washed with PBS with 0.2% Tween20 for 30 min and then PBS alone before blocking in 1% bovine serum albumin and 5% Donkey Serum for 1 hr at room temperature. Incubation with primary anti-RFP (rabbit, 1:1000, 600-401-379, Rockland Immunochemicals, USA) and anti-GFP (chicken, 1:1000, ab13970, Abcam, UK) antibody was for 24 hours at room temperature in blocking solution before washed with PBS with 0.2% Tween20 followed by PBS alone. Incubation with secondary antibodies (1:500, anti-rabbit AlexaFluor594, anti-chicken AlexaFluor488, Invitrogen, UK) was for 1 hr in blocking solution at room temperature. Of note, the usual permeabilization step with Triton X-100 was not used in any of the steps to avoid disruption of the cytoplasmic membrane. Images were acquired using Axioskope2 microscope and Axiovision software (Zeiss, Germany). All images were converted to 8-bit and merged using ImageJ (Fiji).

*Adipose and liver histology quantification*

Adipocyte area was measured in QuPath after manual adipocyte identification using the wand detection tool. Typically this selected entire adipocytes, but some adjustments were required. At least 100 adipocytes were measured per adipose depot, a number sufficient for accurate representation of adipose depots (Parlee *et al.*, 2014). Adipocyte areas from QuPath were binned in GraphPad Prism into 1000µm$^2$ bins. Calculation of adipocyte numbers were based on assumed adipocyte sphericity, and an assumed adipose tissue density of 0.96g/ml (Entenman *et al.*, 1958), with depot masses divided by mean adipocyte mass to calculate total adipocyte number per depot. Lipid content and PSR, SBB, and DAB staining in adipose and liver sections was measured with an automated self-defined pixel classifier.

*Gene Expression Analysis*

Total RNA was extracted from 50 mg samples of liver and adipose depots by TRIzol$^{TM}$ according to the manufacturer's protocol. Concentration of eluted RNA was measured using the NanoDrop ONE before dilution to 100ng/µL in nuclease-free water. Reverse transcription was undertaken using the High Capacity cDNA Reverse Transcription Kit (Applied Biosystems) in the Eppendorf Mastercycler X50s, using 1000ng RNA per reaction. cDNA solutions were diluted 1 in 4. Real-time quantitative PCR (RT-qPCR) was performed using TaqMan reagents including minor groove binder probes (Hein and Bodendorf, 2007) on a LightCycler® 480 Instrument II (Roche). *Gapdh* was evaluated as a housekeeping control gene. Primer efficiency was calculated by standard dilution curve prior to experimental reactions, and reactions were run in triplicate with reverse transcriptase negative and non-template controls run on the same plate. Crossing point (Cp) values were calculated using LightCycler 480 software using the Abs quant/ 2nd derivative max function, and values for *Alms1* (probe Mm01189441_m1) normalised to *Gapdh* values after adjusting for primer efficiency, as described by Pfaffl (Pfaffl, 2001).

*Plasma biochemistry*

Plasma biochemical assays were undertaken by the Core Biochemical Assay Laboratory (CBAL) at the Cambridge University Hospitals NHS Foundation Trust. The Siemens Dimension EXL autoanalyser was used to quantify triglyceride, total cholesterol, HDL-C, alanine transaminase (ALT) and aspartate transaminase (AST). LDL-C is calculated from the triglyceride, HDL-cholesterol and total cholesterol concentrations as described in the Friedwald formula (LDL-C = Total cholesterol – HDL-C - (Triglycerides/2.2). Free fatty acids were quantified by enzymatic colorimetric assay and testosterone was quantified by enzyme-linked immunosorbent assay (ELISA); both reactions were measured on the PerkinElmer Victor-3 Plate Reader. Insulin, leptin, adiponectin and inflammatory cytokines were quantified by electrochemiluminescence immunoassay with measurements read on the MesoScale Discovery Sector s600. The reagent and kit details are detailed in supplementary table 1.
